# Chronic semaglutide treatment reveals stage-dependent changes to feeding behavior and metabolic adaptations in male mice

**DOI:** 10.1101/2025.07.24.666640

**Authors:** Harsh Shah, Julio E. Ayala

**Affiliations:** Department of Molecular Physiology & Biophysics, Vanderbilt University School of Medicine, Nashville, TN, 37232; Vanderbilt Mouse Metabolic Phenotyping Center-Live, Nashville, TN, 37232; Vanderbilt Center for Addiction Research, Nashville, TN, 37223

**Keywords:** Glp1 receptor agonist, body weight, food intake, meal pattern, energy expenditure

## Abstract

Glucagon-like peptide-1 receptor (Glp1r) agonists have transformed obesity treatment, but weight loss responses to these drugs vary widely. Elucidating behavioral and metabolic phenotypes throughout Glp1r agonist treatment could identify mechanisms underlying this response spectrum. We characterized food intake, meal patterns, energy expenditure (EE), and substrate oxidation during chronic semaglutide treatment and post-treatment recovery in obese male mice at room temperature (RT) and thermoneutral temperature (TN). Semaglutide-induced weight loss and post-treatment weight regain were similar at RT and TN. Weight loss was divided into three stages at both temperatures: 1) rapid initial weight loss, 2) slower gradual weight loss, and 3) weight maintenance. Initial weight loss was marked by reduced food intake, smaller and less frequent meals, and increased lipid oxidation. Food intake gradually returned to pre-treatment levels through increased meal frequency, while meal size remained suppressed. Lipid oxidation gradually decreased while carbohydrate oxidation increased. Weight-adjusted EE and locomotor activity increased throughout semaglutide treatment. Mice rapidly regained weight after treatment cessation, and this was associated with increased food intake, meal size and frequency, carbohydrate oxidation, EE, and activity. These findings reveal that semaglutide-induced weight loss and regain after treatment cessation involve dynamic, stage-specific changes in feeding behavior, EE, and substrate oxidation.

**ARTICLE HIGHLIGHTS:**

- Although many studies demonstrate acute behavioral and metabolic effects of glucagon-like peptide-1 receptor (Glp1r) agonists, few have assessed chronic effects of these drugs on these phenotypes.
- We wanted to assess changes to various behavioral and metabolic phenotypes throughout a chronic treatment regimen with semaglutide and post-treatment.
- Weight loss in response to chronic semaglutide treatment can be divided into distinct phases, and each phase is characterized by different effects on food intake, meal patterns, energy expenditure, and substrate oxidation.
- Our findings suggest that differences in behavioral changes and/or metabolic adaptations may underlie the degree of weight loss responsiveness to Glp1r agonists.

---

Several drugs have been approved for treating obesity, but most promoted relatively small, albeit clinically-significant, weight loss (∼5%) and several had side effects that removed them from clinical use [1, 2]. Glucagon-like peptide-1 receptor (Glp1r) agonists have dramatically altered the landscape of obesity therapeutics. Liraglutide (∼5-10% weight loss), semaglutide (∼15% weight loss), and the Glp1r/glucose-dependent insulinotropic polypeptide receptor (GIPr) agonist tirzepatide (∼25% weight loss) show significant promise in reducing rates of obesity [3]. Other multi-peptide compounds using Glp1r agonists as anchors are currently in development as treatments for obesity and its metabolic comorbidities.

Surprisingly little is known about the behavioral and metabolic factors mediating the weight loss effect of Glp1r agonists. Reduced caloric intake via activation of neuronal Glp1r is a key contributor to the weight loss effect of these drugs [4, 5]. However, many studies assessing the effect of Glp1r agonists on food intake in rodent models have focused on acute (24h) timelines, during which nausea/visceral illness effects are predominant [6, 7]. Chronic studies show that reduced food intake in response to Glp1r agonists in rodents is transient [5, 8, 9]. Furthermore, food intake is comprised of several factors such as meal frequency, duration, and size, and rodent studies suggest that changes in these factors can influence body weight even as absolute caloric intake is unaffected [10–16]. Thus, whether and how Glp1r agonists influence meal patterns remains largely unexplored.

Another contributor to changes in body weight is energy expenditure (EE). Studies show that Glp1r agonists either increase [17–19] decrease [20, 21], or do not influence [8, 22–24] EE. These inconsistent findings could be attributed to differences in the duration of treatment and the manner in which rates of EE are presented. A common convention is to normalize rates of EE by body weight or a factor proportional to body weight. Since Glp1r agonists promote weight loss, normalization to a lower body weight can artificially, and possibly incorrectly, conclude that these drugs increase EE. Furthermore, since all the weight loss does not occur at once, normalizing EE to body weight in a group with fluctuating body weight can be problematic.

We address these gaps in knowledge by investigating the effect of Glp1r agonist treatment in obese male mice on food intake, meal patterns, and body weight-adjusted EE throughout a chronic dosing regimen and following cessation of treatment. Most individuals regain a significant amount of weight after stopping Glp1r agonist treatment [25–27], so investigating the behaviors and metabolic factors driving this phenomenon is translationally relevant. We demonstrate that the change in body weight in response to Glp1r agonist treatment occurs in three sequential stages characterized by 1. rapid weight loss, 2. gradual weight loss, and 3. weight maintenance. In support of previous studies, we show that Glp1r agonist treatment reduces food intake, but food intake returns to pre-treatment levels during the gradual weight loss phase. Each phase is characterized by different effects on meal patterns and substrate oxidation. Furthermore, we used daily body weight-adjusted EE using analysis of covariance (ANCOVA) and Glp1r agonist treatment at thermoneutral temperature to measure effects on EE. We also demonstrate that specific changes to meal patterns and EE can potentially contribute to the rapid weight regain observed in mice upon cessation of Glp1r agonist treatment. Altogether, these studies form the basis for future investigations into molecular mechanisms that regulate specific feeding behaviors and EE that contribute to the weight loss effects of Glp1r agonists.

## RESEARCH DESIGN AND METHODS

### Animals

8-12 week-old male C57Bl/6J mice were fed a 60% high fat diet (HFD: 5.24 kcal/g; D12492, Research Diets Inc.) for 9-12 weeks to induce obesity at a housing temperature of ∼22°C. Mice were maintained on a 12- to 12-h light-dark cycle (0600-1800) with ad-libitum access to the food and water. Experimental procedures were approved by the Institutional Animal Care and Use Committee at Vanderbilt University.

### Chronic Glp1r Agonist Dosing Experiments

Following onset of obesity, one cohort of mice remained at room temperature (RT, 22°C), while the other was acclimated to a thermoneutral (TN) temperature of 29°C for 1 week prior to transfer to metabolic chambers. Mice were single-housed and transferred to Promethion Metabolic chambers (Sable Systems International) at the Vanderbilt Mouse Metabolic Phenotyping Center (VMMPC) and acclimated for 3 days (for mice at RT) to 6 days (for mice at TN). Mice were then treated with either Vehicle (Saline) or semaglutide (Novo Nordisk) subcutaneously (n=7-8 per group, 60 µg·kg^-1^·day^-1^, 6µL/g of BW) for 21 days, followed by a 7- day washout period. Mice had ad-libitum access to 60% HFD and water.

### Metabolic Measurements

Food intake, meal patterns, water intake, locomotor activity, respiratory exchange ratio, and energy expenditure (EE) were continuously measured across five-minute intervals. Total daily EE was calculated as the average EE (kcal/hr) each day and multiplied by 24. Analysis of covariance (ANCOVA) was performed using the Mouse Metabolic Phenotyping Center (MMPC) EE Analysis web tool (https://mmpc.org/shared/regression.aspx) with daily body weight used as a covariate. Activity was measured by counting infrared beam breaks in the X, Y, and Z plane and multiplying by the distance between beams. Carbohydrate and lipid oxidation rates were derived from Frayn’s equation [28] and protein oxidation was assumed to be zero. Body composition (NMR, Bruker Optics) was measured prior to vehicle/Glp1r agonist treatment, after the last day of treatment, and at day 6-7 following cessation of treatment.

### Meal Parameters

A meal was defined as food removal spanning at least 30 seconds, a minimum of 10 milligrams to account for inherent sensor limitations, a maximum of 500 milligrams to account for spillage, and a threshold of 5 minutes for inter-meal intervals based on prior rodent literature. A satiety index was calculated as the ratio of intermeal intervals to meal sizes. 60% HFD was packed in the hopper to reduce spillage. Cages were inspected daily for excessive spillage (food on cage bottom) and noted. If total food intake and meal number for a given day was greater than 1.5x standard deviation of the mean for its respective feeding period, then this was considered excessive spillage, and intake and meal pattern data for that day were excluded.

### Statistical Analysis

Raw data were processed using the Sable Systems Macro Interpreter (v23.6). Graphs were generated and data statistically analyzed using repeated measures two-way ANOVA or mixed-effect analysis with Tukey’s multiple comparison test using GraphPad Prism (v10). Significance was set at p<0.05.

## RESULTS

### Semaglutide-induced weight loss is characterized by three distinct phases

Following acclimation to metabolic chambers, mice were treated with vehicle or semaglutide for 21 days followed by a 7-day recovery period (**Figure 1A**). Mice housed at room temperature (RT) displayed weight loss characterized by three phases in terms of absolute body weight (**Figure 1B**) and percent change in body weight (**Figure 1C**). Phase 1 (Day 0-7) was characterized by rapid weight loss, particularly following the first day of treatment. Phase 2 (Day 7-14) was characterized by a gradual decrease in body weight. Phase 3 (Day 14-21) was characterized by reaching a plateau of weight loss and a weight maintenance phase. This same pattern was observed in mice housed at TN (**Figure 1D-E**). Regardless of ambient temperature, all mice rapidly began to regain body weight upon cessation of semaglutide treatment (**Figure 1B-E**). Semaglutide-induced weight loss was primarily driven by loss of fat mass with a smaller contribution of fat free mass regardless of temperature (**Figure 1F-I**).

**Fig 1.**
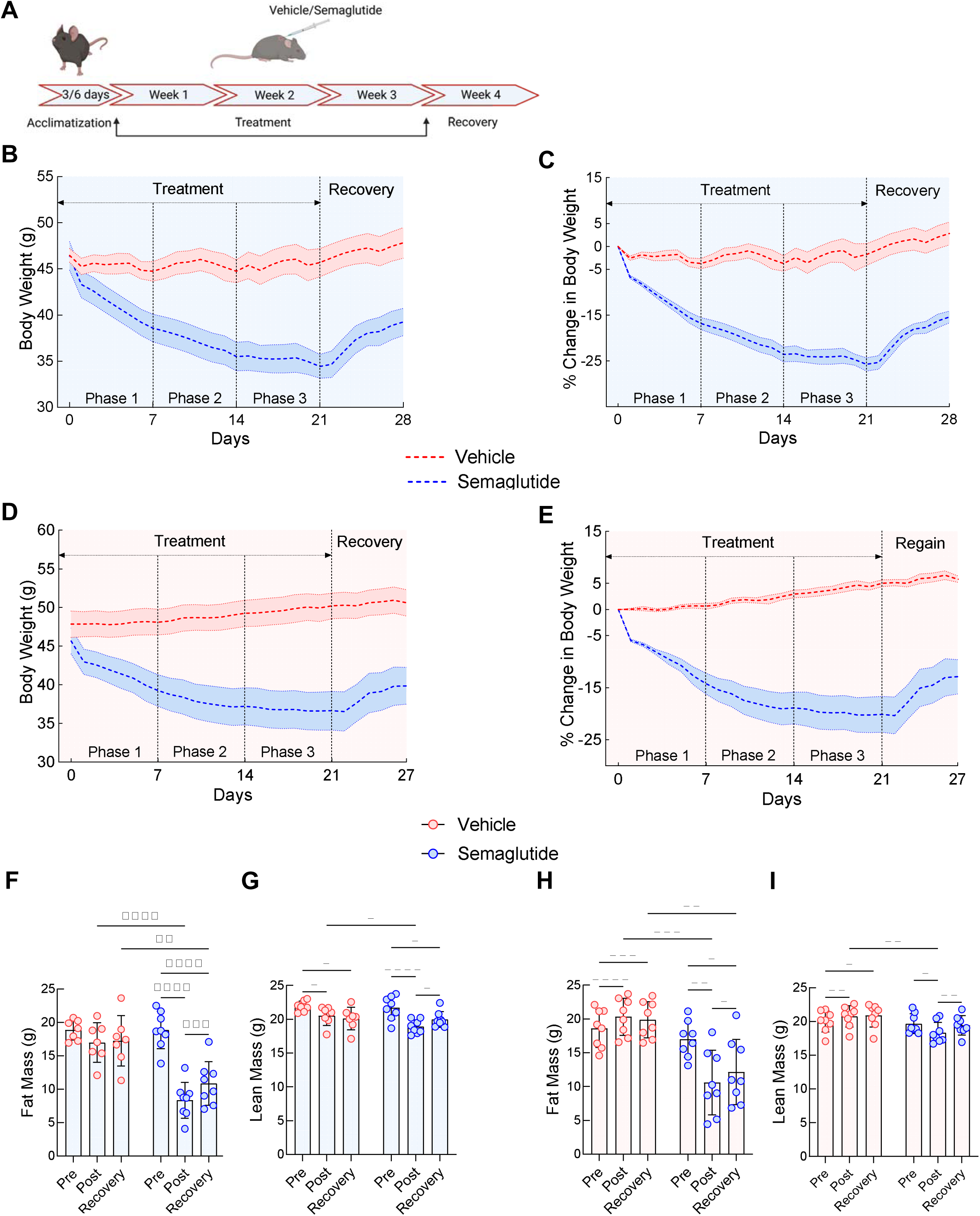
Effect of semaglutide treatment on weight loss and body composition room temperature (RT, 22 °C) and thermoneutral temperature (TN, 29 °C). **A.** Study overview, male diet-induced obese mice were acclimatized for 3 days (RT) and 6 days (TN) in metabolic chambers followed by saline or semaglutide (60 µg·kg^-1^·day^-1^, 6µL/g of volume) treatment for three weeks and no treatment for one week. **B.** Absolute body weight (Time x Treatment - F (2.443, 31.76) = 34.47, P<0.0001) and **C.** percent change in body weight at RT (Time x Treatment - F (2.492, 32.40) = 35.52, P<0.0001). **D.** Absolute body weight (Time x Treatment - F (1.731, 24.23) = 22.50, P<0.0001) and **E.** percent change in body weight at TN (Time x Treatment - F (1.740, 24.37) = 20.81, P<0.0001). **F.** Fat mass (Time x Treatment - F (1.339, 17.41) = 43.98, P<0.0001) and **G.** lean mass (Time x Treatment - F (1.535, 19.96) = 4.102, P<0.0001) pre-treatment (Pre), post-treatment (Post) and at the end of the recovery period (Recovery) at RT. **H.** Fat mass (Time x Treatment - F (1.224, 17.14) = 34.17, P<0.0001) and **I.** lean mass (Time x Treatment - F (1.790, 25.06) = 14.50, P<0.001) pre-treatment (Pre), post-treatment (Post), and at the end of the recovery period (Recovery) at TN. Data are shown as mean ± SEM for *N*=7-8/group. * p< 0.05, ** p<0.01, *** p<0.001, **** p<0.0001.

### Food intake and meal patterns change in a stage-dependent manner in response to chronic semaglutide

Food intake decreased dramatically following the first semaglutide treatment (**Figure 2A**) and remained lower during the rapid weight loss phase, but it gradually increased to levels comparable to those observed in vehicle treated mice by the weight maintenance phase (**Figure 2A-B**). Immediately after cessation of semaglutide treatment, food intake significantly increased to levels above those observed in vehicle-treated mice (**Figure 2A-B**).

**Fig 2.**
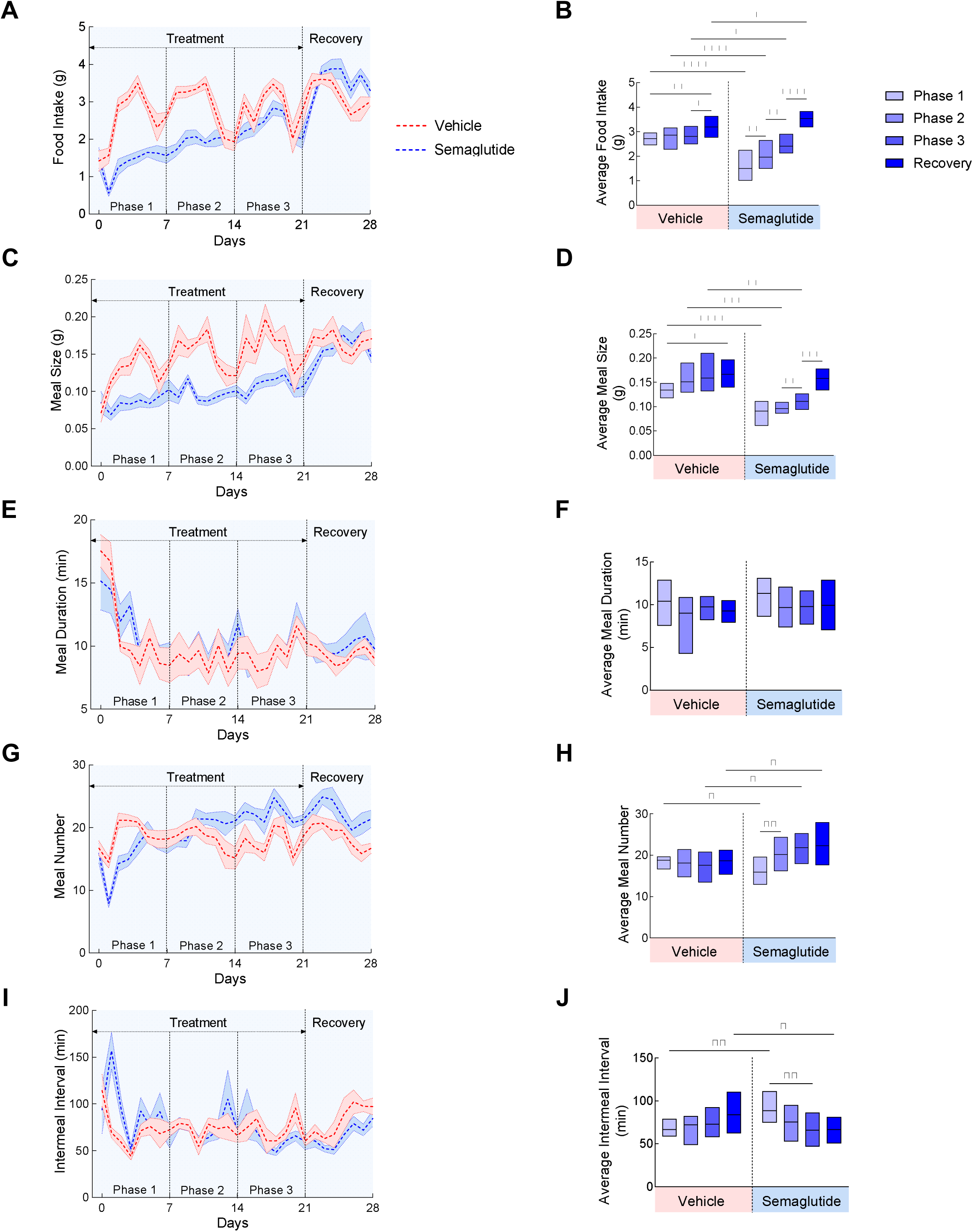
Effect of semaglutide treatment on food intake and meal patterns at RT. **A.** Daily food intake (Time x Treatment (F (7.366, 95.76) = 8.140, p<0.0001) and **B.** average food intake for each phase (Time x Treatment - F (3, 39) = 27.05, p<0.0001). **C** Daily meal size (Time x Treatment - F (6.456, 83.93) = 3.570, p<0.01) and **D.** average meal size for each phase (Time x Treatment - F (3, 39) = 7.701, p<0.001). **E.** Daily meal duration (Time x Treatment - F (6.316, 82.11) = 0.6408, p>0.05) and **F.** average meal duration for each phase (Time x Treatment - F (3, 52) = 0.2188, p>0.05). **G.** Daily meal number (Time x Treatment - F (8.622, 112.1) = 5.057, p<0.001) and **H.** average meal number for each phase (Time x Treatment - F (3, 39) = 20.52, p<0.001). **I.** Daily intermeal interval (Time x Treatment - F (6.460, 83.97) = 2.096, p=0.057) and **J.** average intermeal interval for each phase (Time x Treatment - F (3, 39) = 8.645, p<0.001). Data are shown as mean ± SEM for *N*=7-8/group. * p< 0.05, ** p<0.01, *** p<0.001, **** p<0.0001.

Decreased food intake in semaglutide-treated mice was associated with decreased meal size (**Figure 2C-D**). Although meal size gradually increased throughout the treatment period, it did not reach levels equivalent to those observed in vehicle-treated mice. Cessation of treatment resulted in a rapid increase in meal size (**Figure 2C-D**) that paralleled the increased food intake. Meal duration was unaffected by semaglutide treatment (**Figure 2E-F**). Interestingly, meal numbers decreased following the first day of semaglutide treatment, but they rapidly increased and reached levels above those observed in vehicle-treated mice by the weight maintenance phase and remained elevated after ending treatment (**Figure 2G-H**). Intermeal interval rapidly increased following the first day of semaglutide treatment but then decreased (**Figure 2I-J**). Since meal size is indicative of satiation, these findings suggest that enhanced satiation significantly contributes to semaglutide-induced weight loss, particularly since the biggest differences compared to vehicle-treated mice were observed during the rapid and gradual weight loss phases. Conversely, satiety, which contributes to meal numbers, initially increases but rapidly deteriorates (**Supplemental Figure 1A-B**).

### Chronic semaglutide treatment increases body weight-adjusted energy expenditure and locomotor activity

Absolute EE decreased rapidly following the first semaglutide treatment and gradually continued to decrease below what is observed in vehicle-treated mice (**Figure 3A-B**). We performed analysis of covariance (ANCOVA) to account for daily body weight as a covariate for EE. Contrasting the result from the absolute EE, following an initial drop after the first semaglutide treatment, EE remained steady and higher relative to vehicle-treated mice when accounting for changes in body weight (**Figure 3C-D**). These differences in EE were consistent with a relatively higher level of locomotor activity in semaglutide-treated mice compared to vehicle-treated mice, particularly during the weight maintenance phase (**Figure 3E-F**).

**Fig 3.**
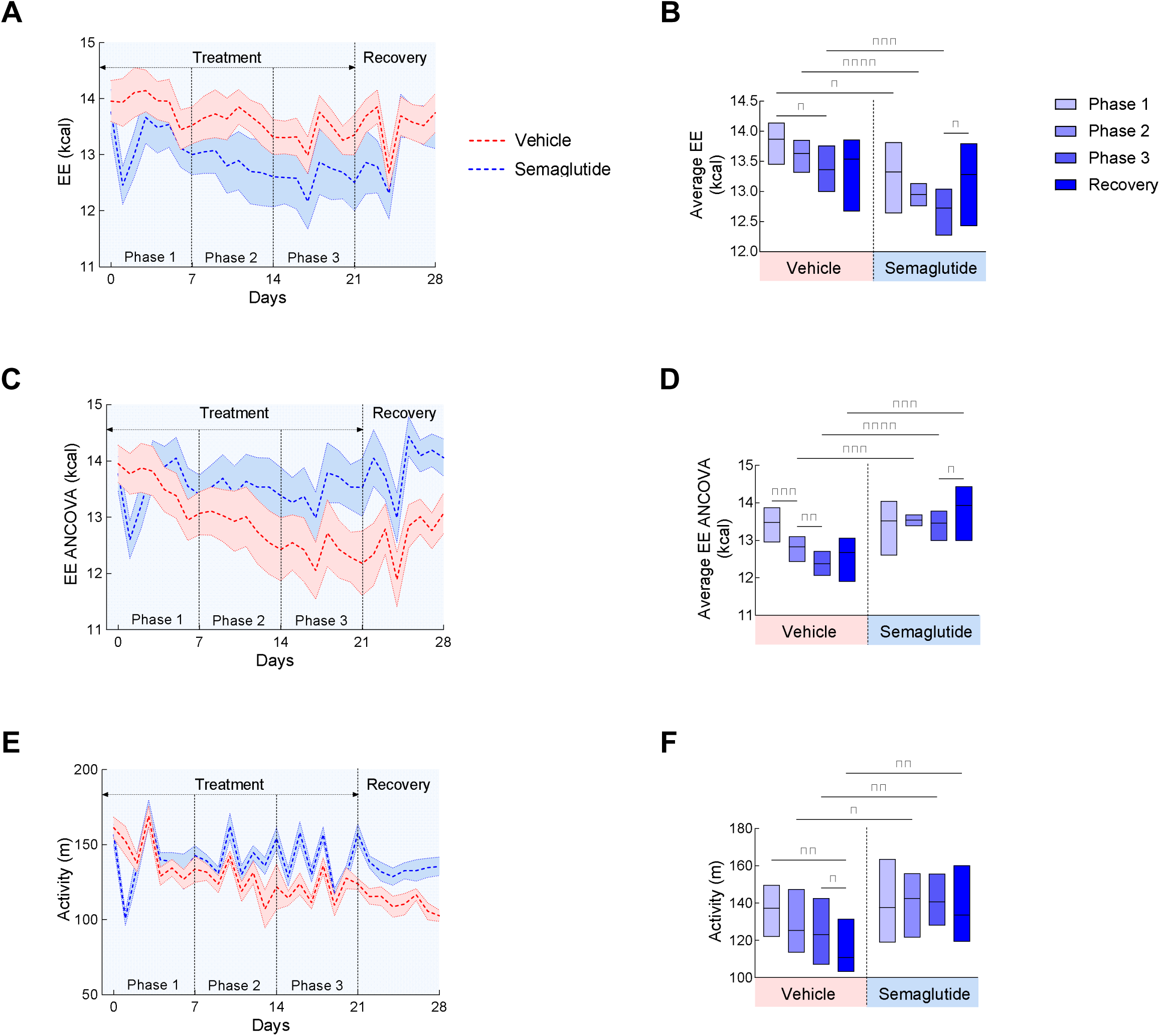
Effect of semaglutide treatment on energy expenditure at RT. **A.** Daily energy expenditure (EE) (Time x Treatment - F (28, 364) = 2.151, p<0.001) and **B.** average EE for each phase (Time x Treatment - F (3, 36) = 1.370, p>0.05). **C.** Daily analysis of covariance (ANCOVA)-adjusted EE with body weight as a covariate and **D.** average ANCOVA-adjusted EE for each phase (Time x Treatment - F (3, 36) = 9.178, p<0.001). **E.** Daily activity (Time x Treatment - F (6.273, 81.55) = 4.267, p<0.001) and **F.** average activity in each phase (Time x Treatment - F (3, 39) = 3.802, p<0.05). Data are shown as mean ± SEM for *N*=7-8/group. * p< 0.05, ** p<0.01, *** p<0.001, **** p<0.0001.

### Chronic semaglutide treatment promotes changes in substrate oxidation

Consistent with the composition of the 60% HFD and feeding patterns, the respiratory exchange ratio (RER) in vehicle-treated mice vacillated steadily between 0.75 and 0.8, indicative of greater lipid oxidation relative to carbohydrate oxidation (**Figure 4A-B**). This was supported by calculations of lipid and carbohydrate oxidation rates (**Figure 4C-F**). Contrasting this, RER remained lower in semaglutide-treated mice, but it gradually increased to levels similar to those observed in vehicle-treated mice by the weight maintenance phase. This is indicative of a time-dependent shift in substrate oxidation whereby carbohydrate oxidation increased and lipid oxidation decreased throughout semaglutide treatment (**Figure 4C-F**). Stopping semaglutide treatment rapidly increased carbohydrate oxidation and decreased lipid oxidation (**Figure 4C-F**).

**Fig 4.**
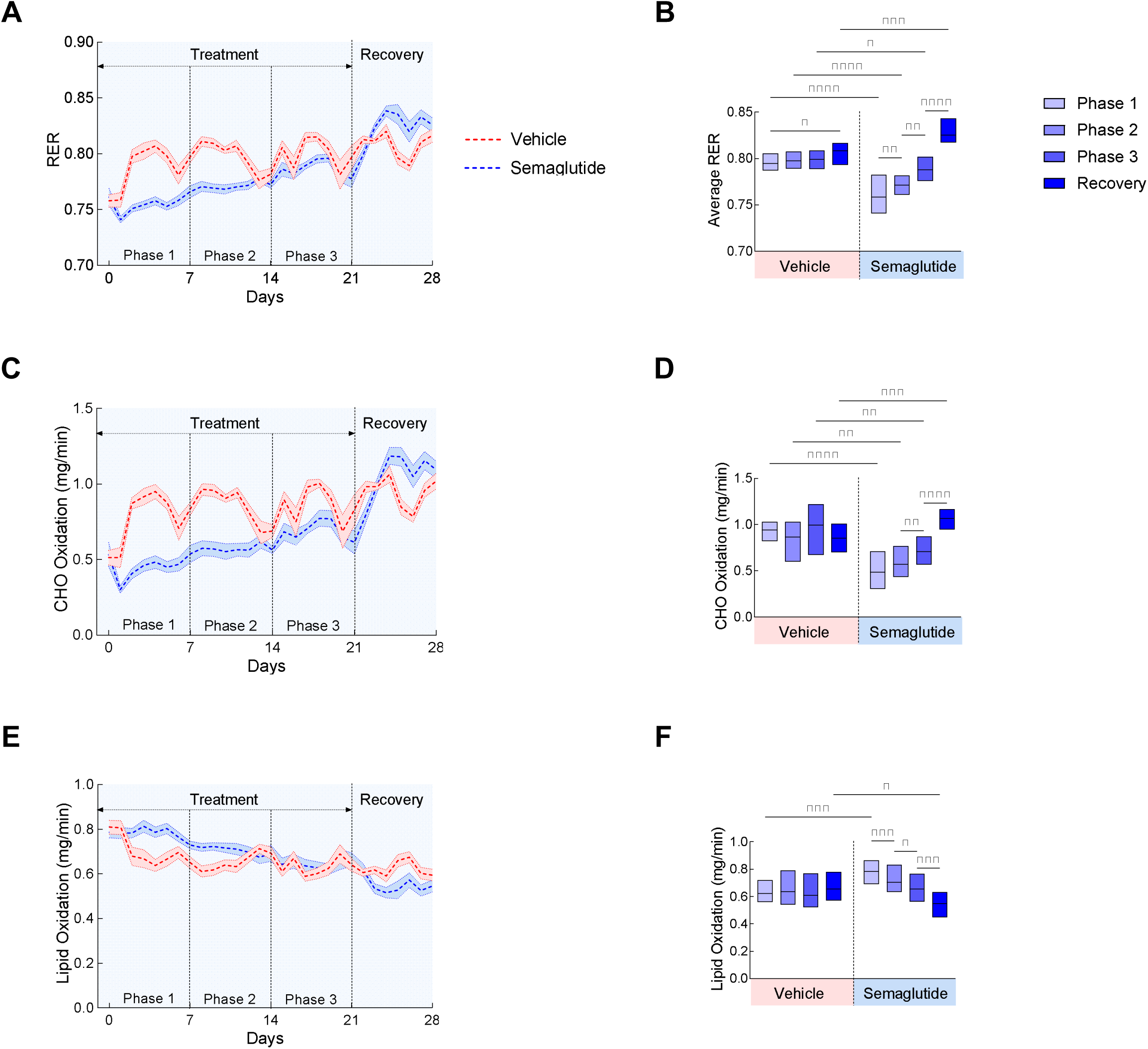
Effect of semaglutide treatment on respiratory exchange ratio (RER) and carbohydrate and lipid oxidation rates at RT. **A.** Daily RER (Time x Treatment - F (6.976, 90.69) = 9.756, p<0.0001) and **B.** average RER for each phase (Time x Treatment - F (3, 39) = 31.87, p<0.0001). **C.** Daily carbohydrate (CHO) oxidation rate (Time x Treatment - F (6.906, 89.78) = 10.45, p<0.0001) and **D.** Average carbohydrate (CHO) oxidation rate for each phase (Time x Treatment - F (3, 39) = 22.29, p<0.0001). **E.** Daily lipid oxidation rate (Time x Treatment - F (5.942, 77.25) = 8.687, p<0.0001) and **F.** lipid oxidation rate for each phase (Time x Treatment - F (3, 39) = 12.91, p<0.0001). Data are shown as mean ± SEM for *N*=7-8/group. * p< 0.05, ** p<0.01, *** p<0.001, **** p<0.0001.

### Thermoneutrality does not affect feeding patterns in response to semaglutide treatment

Similarly to what we observed in mice housed at RT, semaglutide treatment initially reduced food intake and meal size in mice housed at TN, but this gradually increased to levels similar to what was observed in vehicle-treated mice by the weight maintenance phase (**Figure 5A-D**). Cessation of semaglutide treatment significantly increased food intake and meal size (**Figure 5A-D**). Meal duration was unchanged and relatively equivalent in vehicle- vs. semaglutide-treated mice, although meal duration increased after stopping semaglutide treatment (**Figure 5E-F**). Following a rapid decline after initiating semaglutide treatment, meal numbers gradually increased to reach equal levels to those observed in vehicle-treated mice, but unlike mice housed at RT, meal numbers did not increase above those observed in vehicle treated mice in mice at TN (**Figure 5G-H**). As expected, intermeal interval initially increased upon initiation of semaglutide treatment and subsequently decreased (**Figure 5I-J**). Similarly to mice housed at RT, this shows that satiety initially increases but rapidly decreases with semaglutide treatment in mice at TN (**Supplemental Figure 1C-D**).

**Fig 5.**
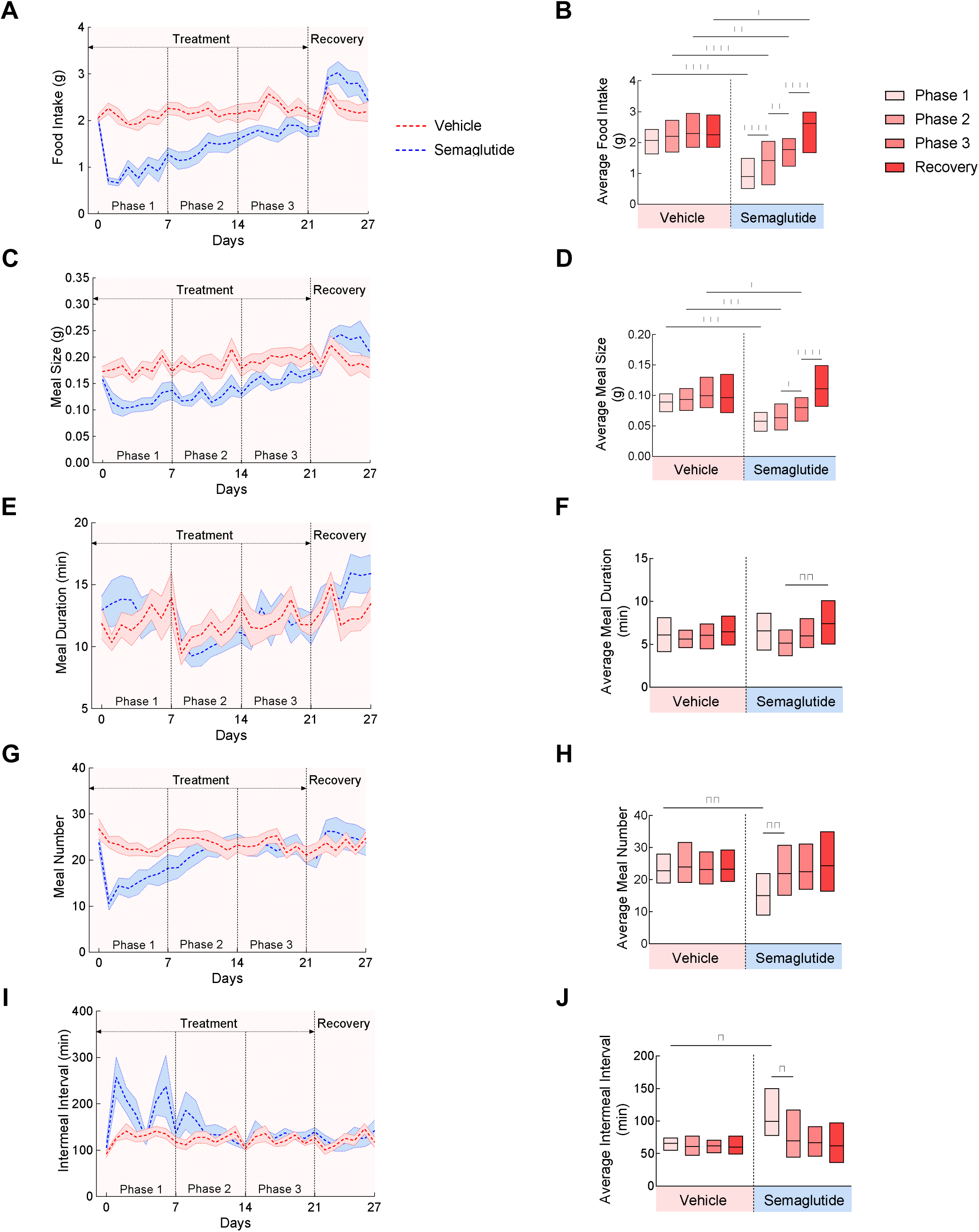
Effect of semaglutide treatment on food intake and meal patterns at TN. **A.** Daily food intake (Time x Treatment - F (6.695, 93.73) = 13.06, p<0.0001) and **B.** average food intake for each phase (Time x Treatment - F (3, 42) = 46.06, p<0.0001). **C.** Daily meal size (Time x Treatment - F (7.187, 100.4) = 4.201, p<0.001) and **D.** average meal size for each phase (Time x Treatment - F (3, 42) = 14.42, p<0.0001). **E.** Daily meal duration (Time x Treatment - F (7.851, 109.9) = 1.516, p>0.05) and **F.** Average meal duration for each phase (Time x Treatment - F (3, 28) = 1.623, p>0.05). **G.** Daily meal number (Time x Treatment - F (6.237, 87.31) = 5.027, p<0.001) and **H.** average meal number for each phase (Time x Treatment - F (3, 42) = 10.35, p<0.001). **I.** Daily intermeal Interval (Time x Treatment - F (4.812, 67.19) = 3.277, p<0.05) and **J.** average intermeal interval for each phase (Time x Treatment - F (3, 42) = 4.260, p<0.05). Data are shown as mean ± SEM for *N*=8/group. * p< 0.05, ** p<0.01, *** p<0.001, **** p<0.0001.

### EE increases in response to semaglutide treatment at TN

Absolute EE initially decreased upon semaglutide treatment, but it rapidly rebounded to levels similar to those observed in vehicle-treated mice (**Figure 6A-B**). Accounting for changes in body weight shows that, similarly to what was observed in mice at RT, EE was higher in semaglutide-treated mice throughout the treatment period, and this was further increased upon stopping treatment (**Figure 6C-D**). This increase in EE was accompanied by increased locomotor activity (**Figure 6E-F**). By incorporating the contribution of daily changes in body weight and thermoneutrality, we demonstrate that an increase in EE likely contributes to the weight loss effect of semaglutide.

**Fig 6.**
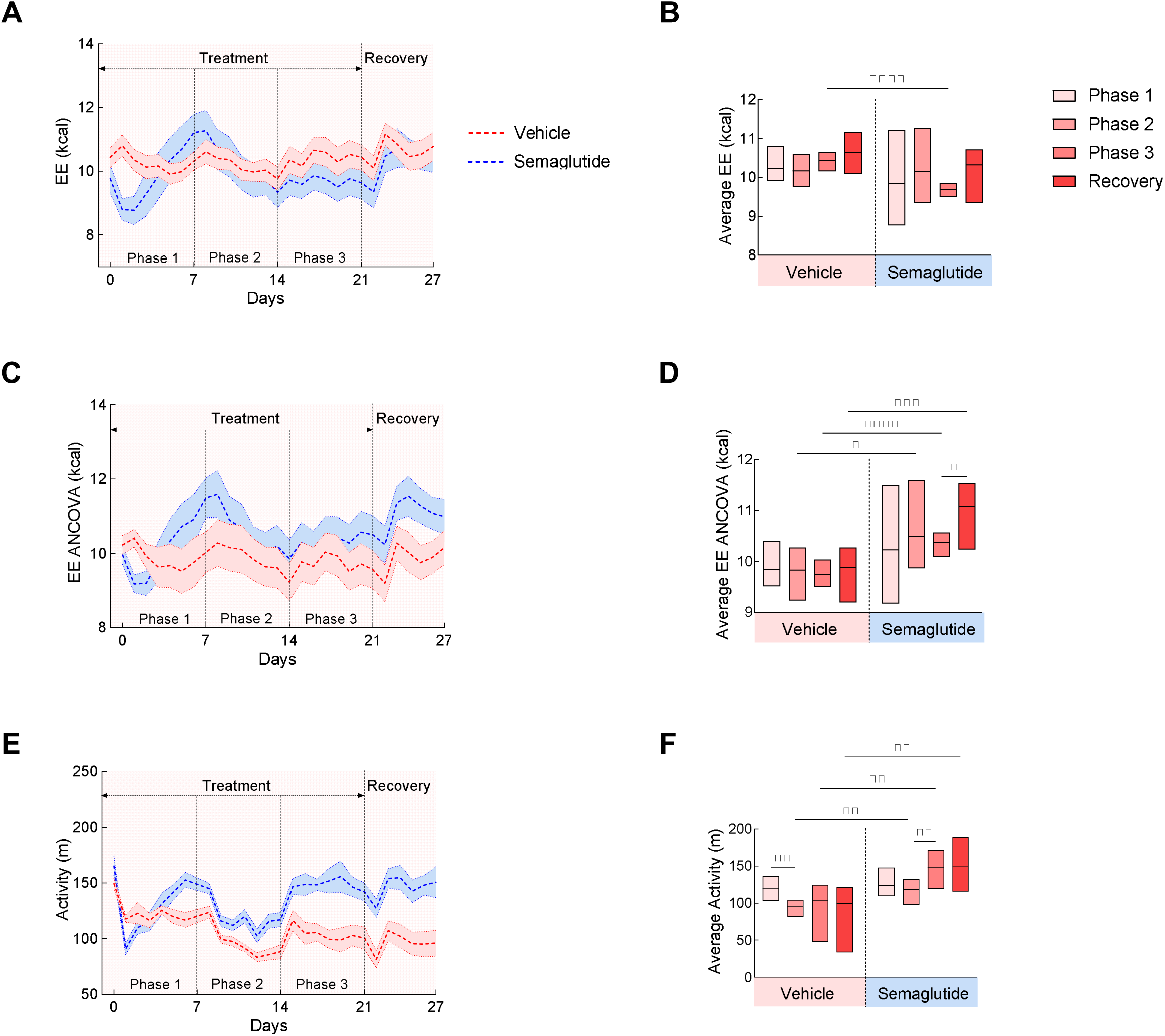
Effect of semaglutide treatment on energy expenditure at TN. **A.** Daily energy expenditure (EE) (Time x Treatment - F (2.417, 33.84) = 6.134, p<0.01) and **B.** average EE for each phase (Time x Treatment - F (3, 46) = 1.316, p>0.05). **C.** Daily analysis of covariance (ANCOVA)-adjusted EE with body weight as a covariate and **D.** average ANCOVA-adjusted EE for each phase (Time x Treatment - F (3, 46) = 1.599, p>0.05). **E.** Daily activity (Time x Treatment - F (4.664, 65.30) = 5.856, p<0.001) and **F.** average activity for each phase (Time x Treatment - F (1.624, 22.74) = 10.80, p<0.001). Data are shown as mean ± SEM for *N*=8/group. * p< 0.05, ** p<0.01, *** p<0.001, **** p<0.0001.

### Chronic semaglutide treatment promotes similar changes in substrate oxidation at TN compared to RT

RER in vehicle-treated mice remained constant at ∼0.75 (**Figure 7A-B**) consistent with greater lipid relative to carbohydrate oxidation (**Figure 7C-F**). RER decreased upon initiation of semaglutide treatment and remained lower throughout the rapid weight loss period, but it gradually increased during the slow weight loss phase and reached levels similar to those observed in vehicle-treated mice by the weight maintenance phase (**Figure 7A-B**). Similarly to mice treated at RT, this is indicative of an initial increase in lipid oxidation followed by an increase in carbohydrate oxidation (**Figure 7C-F**). However, in mice at TN, elevated lipid oxidation occurred later during the transition from the rapid to the gradual weight loss period and abruptly ended during the transition into the weight maintenance period, whereas carbohydrate oxidation was significantly lower during the rapid weight loss period and increased during the gradual weight loss period (**Figure 7C-F**). Similarly to mice treated at RT, cessation of semaglutide treatment rapidly increased carbohydrate oxidation and rapidly decreased lipid oxidation (**Figure 7C-F**).

**Fig 7.**
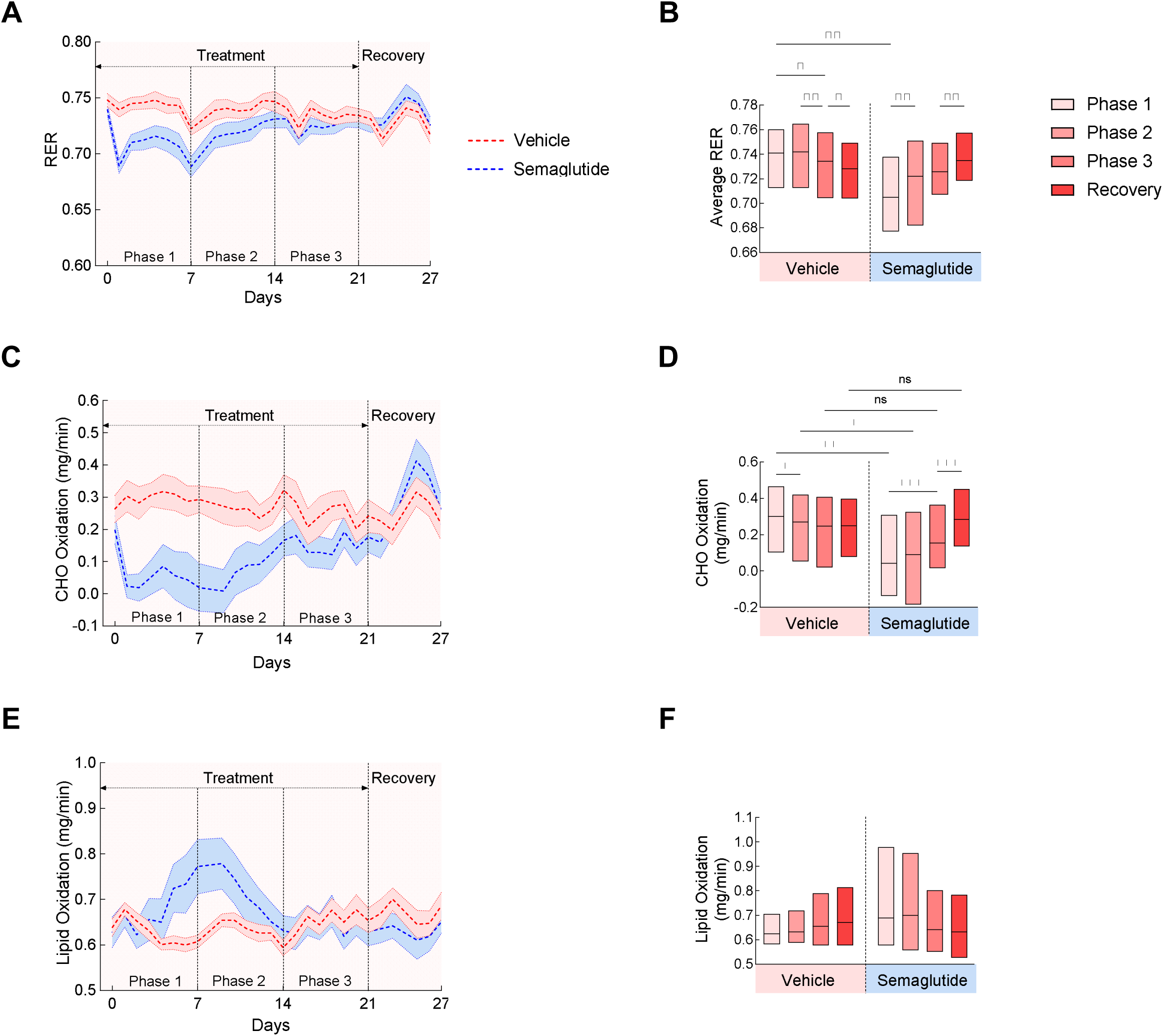
Effect of semaglutide treatment on respiratory exchange ratio (RER) and carbohydrate and lipid oxidation rates at TN. **A.** Daily RER (Time x Treatment - F (6.683, 93.56) = 10.87, p<0.0001) and **B.** average RER for each phase (Time x Treatment - F (3, 42) = 40.67, p<0.0001). **C.** Daily carbohydrate (CHO) oxidation rate (Time x Treatment - F (5.543, 77.61) = 14.69, p<0.0001) and **D.** average carbohydrate (CHO) oxidation rate for each phase (Time x Treatment F (3, 42) = 49.61, p<0.0001). **E.** Daily lipid oxidation rate (Time x Treatment - F (2.549, 35.68) = 6.666, p<0.01) and **F.** average lipid oxidation rate for each phase (Time x Treatment - F (3, 42) = 6.584, p<0.01). Data are shown as mean ± SEM for *N*=8/group. * p< 0.05, ** p<0.01, *** p<0.001, **** p<0.0001.

## DISCUSSION

The degree of weight loss achieved by the Glp1r agonist semaglutide and the dual Glp1r/GIPr agonist tirzepatide has not been observed with previous obesity therapeutics, and new Glp1r agonist-based drugs will likely rival bariatric surgery for degree of weight loss. Nevertheless, not all patients display equal degrees of weight loss, and some are completely unresponsive [29, 30]. This prompts a comprehensive investigation into the behavioral and metabolic changes induced by Glp1r agonists with the goal of revealing factors that contribute to the variability in the response to these drugs. The present studies demonstrate that semaglutide-induced weight loss can be divided into different stages based on the rate of weight loss, and these stages are characterized by different feeding behaviors and metabolic parameters. Initial rapid weight loss results from decreased food intake, smaller and less frequent meals, and a reliance on fat oxidation. As weight loss gradually slows and reaches a nadir, meal sizes remain small, yet food intake returns to pre-treatment levels due to increased meal frequency. Furthermore, reliance on lipid oxidation decreases whereas carbohydrate oxidation increases. Weight adjusted EE remains elevated in response to chronic semaglutide. These findings suggest that differences in factors that regulate meal patterns as well as metabolic adaptations may contribute to the different responsiveness to these drugs in individuals with obesity.

Although weight loss in response to Glp1r agonists is attributed to decreased caloric intake, this effect is not a permanent feature of chronic treatment with these drugs. Our findings confirm previous studies showing that caloric intake returns to pre-treatment levels after ∼1 week of Glp1r agonist treatment [5, 8, 9]. This coincides with the transition from rapid to gradual weight loss and weight maintenance. A return to pre-treatment caloric intake could result from homeostatic mechanisms being engaged to prevent excessive weight loss and establish a new “setpoint” body weight. Although food intake returns to pre-treatment levels, these intake levels can still be interpreted as being inappropriately low relative to the new, lower body weight. Indeed, upon cessation of treatment, food intake rapidly increases to return body weight to pre-treatment levels. Thus, when accounting for the decreased body weight, semaglutide treatment does maintain caloric intake in a relatively suppressed state.

Assessing meal patterns provides a more comprehensive analysis of factors that contribute to changes in body weight than focusing on absolute caloric intake alone. Meal size is a strong predictor of weight gain in rats switched from low to high fat diets, and genetic mouse models of obesity demonstrate a correlation between weight gain and meal size beyond daily caloric intake [10–16]. Here we show that meal size rapidly decreases in response to semaglutide treatment, and although it gradually increases throughout the treatment period, meal size remains below what is observed in vehicle-treated mice. This is indicative of increased satiation or the feeling of fullness within a meal. Studies in humans suggest that Glp1r agonists reduce food intake in part by increasing satiation [21, 31–33]. This is consistent with mouse studies showing increased neuronal activity in brain regions associated with satiation and meal termination in response to Glp1r agonist treatment [8, 34], although this is an indirect measure and was only assessed in response to an acute treatment. Contrasting the effects on satiation, meal numbers rapidly increase in response to semaglutide treatment and reach levels above what is observed in vehicle-treated mice. This strongly suggests that a long-term effect of semaglutide is to enhance satiation (meal size) but not satiety (meal number). Interestingly, studies in humans suggest that Glp1r agonists enhance satiety as well as satiation [32, 35–39]. This may be indicative of a difference in the response to Glp1r agonists between mice and humans. However, it is important to note that studies showing enhanced satiety in humans were conducted following either acute or relatively short-term Glp1r agonist treatment while the study subjects were still losing weight. Since we observe that enhanced satiety in mice is observed primarily during the weight loss phase of semaglutide treatment, this suggests that increased satiety contributes only to Glp1r agonist-induced weight loss but not weight maintenance.

One potential implication from these findings is that differences in regulating satiation and/or satiety may contribute to the spectrum of Glp1r agonist-induced weight loss in clinical settings. In humans, higher body weight and waist circumference is associated with impaired satiation indicated by higher calories consumed before terminating a meal [40]. Interestingly, this was also associated with greater weight loss in response to another weight loss drug, phentermine-topiramate, although it is not clear whether the correlation to weight loss is with the starting body weight or the impaired satiation [40]. A follow-up study confirmed that individuals with inherently higher calories consumed to reach satiation (CTS), that is impaired satiation, prior to treatment displayed greater weight loss in response to phentermine-topiramate. Surprisingly, the same study showed that lower inherent CTS (i.e., enhanced satiation) prior to treatment was associated with greater weight loss in response to liraglutide [41]. One limitation of this study is that potential changes in satiation during the treatment were not assessed. Nevertheless, these findings suggest that factors that regulate satiation may contribute to the effectiveness of weight loss drugs.

It has been difficult to ascertain the effect of Glp1r agonists on EE based on factors such as the duration of treatment when EE measurements were made as well as the fact that EE is often normalized to body weight, a variable in Glp1r agonist weight loss studies. Here we performed ANCOVA with body weight as a covariate and show that following an initial decrease during the first 24h of semaglutide treatment, EE is restored and remains slightly elevated compared to vehicle treatment. Locomotor activity followed a similar pattern, suggesting that this may be a contributor to the difference in EE between vehicle- and semaglutide-treated mice. This contrasts findings from another study showing that ANCOVA-adjusted EE is not different between vehicle- and semaglutide-treated mice with lean mass being used as a covariate [8]. However, those studies used a lower dose of semaglutide (40 vs. 60 µg·kg^-1^·day^-1^) which did not cause a loss of lean mass. We also observed increased EE and locomotor activity in response to semaglutide in mice maintained at TN, a treatment that is expected to decrease overall basal metabolic rates. These findings suggest that increased EE relative to the change in body weight contributes to the weight loss and weight maintenance effect of semaglutide.

We provide additional insight into potential mechanisms contributing to the weight loss effect of semaglutide by demonstrating changes in substrate oxidation during the treatment period. As expected, vehicle-treated HFD-fed mice primarily rely on lipid oxidation for their energy needs. In agreement with previous findings [8], semaglutide treatment promotes greater lipid oxidation as evidenced by a decrease in RER and an increase in the rates of lipid oxidation. However, as the treatment continues, an interesting pattern emerges whereby lipid oxidation gradually decreases and carbohydrate oxidation gradually increases. One potential explanation is that higher lipid oxidation drives the initial weight loss in response to semaglutide, but the reliance on lipid oxidation wanes as a means to protect remaining fat stores and preventing additional weight loss. A similar pattern emerges in mice treated with semaglutide at TN, although with some key differences. Whereas lipid oxidation rapidly increases upon initiation of semaglutide treatment and then gradually decreases in mice at RT, in mice at TN lipid oxidation dramatically increases during the transition from the rapid to the gradual weight loss phase. Once mice reach the weight maintenance phase, lipid oxidation is not different between vehicle- and semaglutide-treated mice. In mice at TN, carbohydrate oxidation rapidly decreases upon initiation of semaglutide treatment, and then it gradually increases to similar levels as in vehicle-treated mice beginning with the transition to the gradual weight loss phase. Similar patterns of lipid and carbohydrate oxidation have been recently reported in mice and humans treated with tirzepatide [42]. These findings further support the hypothesis that increased lipid oxidation contributes to the weight loss induced by Glp1r agonists. However, our findings also suggest that once peak weight loss is achieved, there is a relative decrease in lipid oxidation and an increase in carbohydrate oxidation that likely prevents additional loss of fat mass.

Cessation of treatment with Glp1r agonists is associated with weight regain in the majority of patients [25–27]. We show that post-recovery weight gain is associated with increased food intake driven by increased meal sizes and a maintenance of higher meal numbers. Interestingly, the relatively higher EE and activity levels associated with semaglutide treatment are preserved upon cessation of treatment. This is counterintuitive since one would expect EE to decrease as a means to accelerate weight regain. The increased EE could be secondary to the elevated food intake. Indeed, we have previously shown that increased food and caloric intake in mice switched from a chow diet to a HFD is accompanied by a rapid increase in EE [43]. The post-semaglutide treatment period is also characterized by a dramatic increase in carbohydrate oxidation while fat oxidation remains low. The fact that this occurs even as HFD consumption increases suggests a metabolic shift towards recovering fat stores to promote weight regain.

The continued development of Glp1-based therapeutics means that this drug class will likely become the standard of care for obesity in the near future. It is, therefore, important to investigate all behavioral, metabolic, and mechanistic properties affected by these drugs.

Surprisingly, most rodent studies to date have focused on events that occur following a single dose of Glp1r agonists. As in humans, a common response to the initiation of treatment with Glp1r agonists in rodents is nausea/malaise, so behaviors and metabolic changes during the first 24h primarily reflect visceral illness responses. The present studies provide important insight into events that occur in response to chronic treatment with semaglutide and its discontinuation. This forms a foundation for future studies that will investigate changes in molecular mechanisms and neural networks at different stages of Glp1r agonist treatment.

### Limitations

these studies were conducted only in male mice, so future studies need to investigate whether similar behavioral and metabolic changes are observed in female mice. We draw conclusions on satiation and satiety from *ad libitum* meal patterns and not from direct measures of satiation and satiety (e.g., from pre-load experiments). Similarly, rates of lipid and carbohydrate oxidation were derived from indirect calorimetry measurements. Future studies using isotopic tracers are needed to derive direct measurements of fuel oxidation and tissue-specific nutrient fluxes. This is important since one component not included in the present calculations is the contribution of changes to protein synthesis and degradation. Since Glp1r agonist treatment promotes loss of lean mass, direct measurements of protein fluxes will be informative. Lastly, mice in the present studies were given access to only one type of diet which is not representative of the real-life condition of patients. Glp1r agonists shift dietary preferences, so future studies should also investigate the effect of chronic treatment with these drugs on meal patterns and metabolic responses in the presence of different diets.

## ACKNOWLEDGMENTS

We would like to thank Dr. Louise Lantier and Ms. Merrygay James of the Vanderbilt Mouse Metabolic Phenotyping Center-*Live* for their assistance with the metabolic chamber studies.

## Author Contributions

HS designed and performed the experiments; HS and JEA analyzed the results; HS and JEA wrote and edited the manuscript.

## Guarantor’s statement

Dr. Julio E. Ayala is the guarantor of this work and takes responsibility for the integrity and accuracy of the statements made in this article.

---

**Fig S1:**
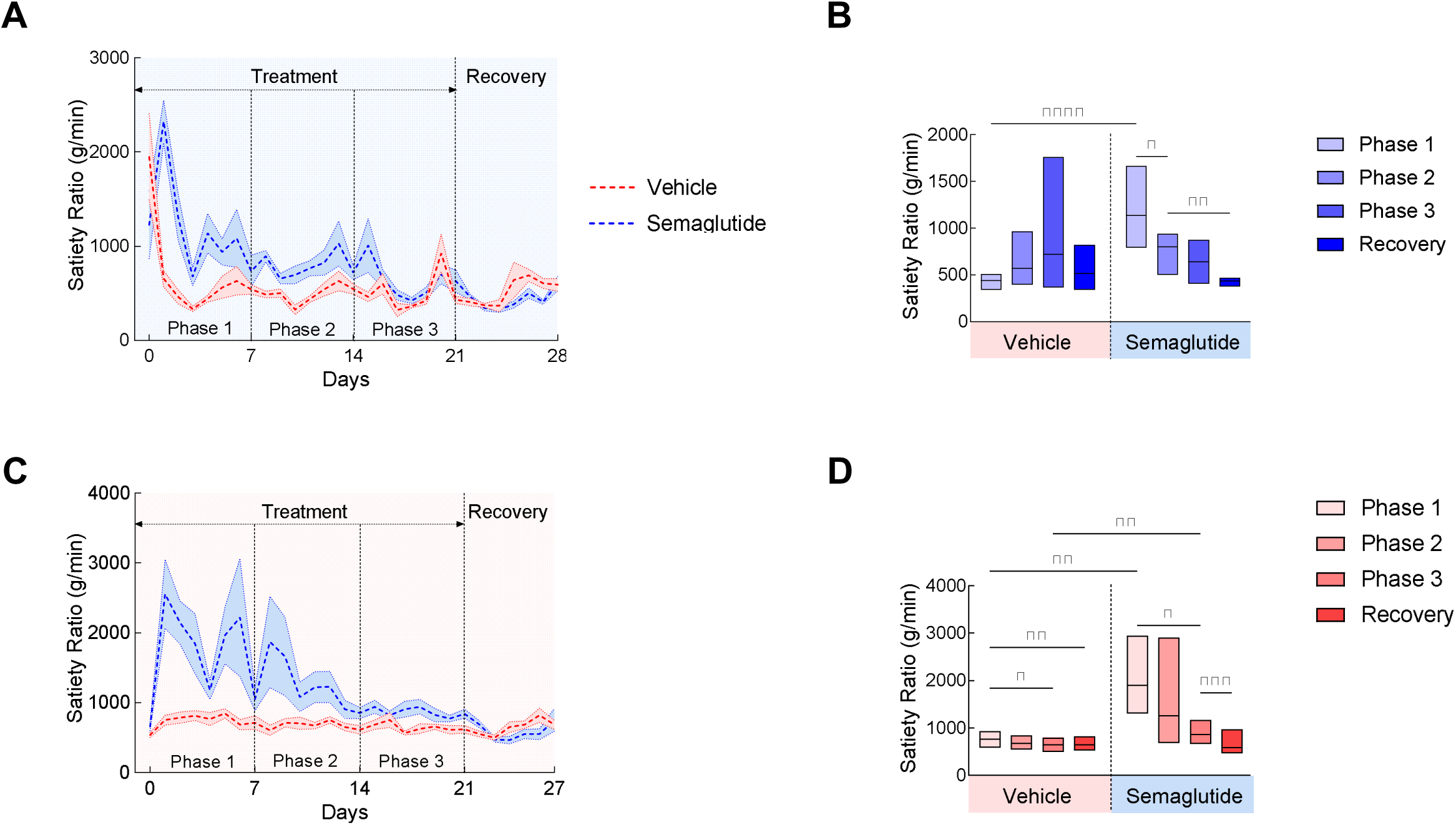
Effect of semaglutide treatment on satiety ratio at RT and TN. **A.** Daily satiety ratio at RT (Time x Treatment - F (4.700, 61.11) = 4.631, p<0.01) and **B.** average satiety ratio for each phase at RT (Time x Treatment - F (3, 39) = 11.62, p<0.0001). **C.** Daily satiety ratio at TN (Time x Treatment - F (3.330, 46.37) = 3.713, p<0.05) and **D.** average satiety ratio for each phase at TN (Time x Treatment - F (3, 42) = 11.26, p<0.0001). Data are shown as mean ± SEM for *N*=7- 8/group. * p< 0.05, ** p<0.01, *** p<0.001, **** p<0.0001.

## REFERENCES

1. Bray, G.A., Obesity: a 100 year perspective. Int J Obes (Lond), 2025. 49(2): p. 159–167.

2. Henderson, K., et al., Effectiveness and safety of drugs for obesity. Bmj, 2024. 384: p. e072686.

3. Drucker, D.J., GLP-1-based therapies for diabetes, obesity and beyond. Nat Rev Drug Discov, 2025.

4. Sisley, S., et al., Neuronal GLP1R mediates liraglutide’s anorectic but not glucose-lowering effect. J Clin Invest, 2014. 124(6): p. 2456–63.

5. Adams, J.M., et al., Liraglutide Modulates Appetite and Body Weight Through Glucagon-Like Peptide 1 Receptor-Expressing Glutamatergic Neurons. Diabetes, 2018. 67(8): p. 1538–1548.

6. Lachey, J.L., et al., The role of central glucagon-like peptide-1 in mediating the effects of visceral illness: differential effects in rats and mice. Endocrinology, 2005. 146(1): p. 458–62.

7. Kanoski, S.E., et al., The role of nausea in food intake and body weight suppression by peripheral GLP-1 receptor agonists, exendin-4 and liraglutide. Neuropharmacology, 2012. 62(5-6): p. 1916–27.

8. Gabery, S., et al., Semaglutide lowers body weight in rodents via distributed neural pathways. JCI Insight, 2020. 5(6).

9. Le, T.D.V., et al., Fibroblast growth factor-21 is required for weight loss induced by the glucagon-like peptide-1 receptor agonist liraglutide in male mice fed high carbohydrate diets. Mol Metab, 2023. 72: p. 101718.

10. Furnes, M.W., C.M. Zhao, and D. Chen, Development of obesity is associated with increased calories per meal rather than per day. A study of high-fat diet-induced obesity in young rats. Obes Surg, 2009. 19(10): p. 1430–8.

11. Treesukosol, Y. and T.H. Moran, Analyses of meal patterns across dietary shifts. Appetite, 2014. 75: p. 21–9.

12. Melhorn, S.J., et al., Acute exposure to a high-fat diet alters meal patterns and body composition. Physiol Behav, 2010. 99(1): p. 33–9.

13. Strohmayer, A.J. and G.P. Smith, The meal pattern of genetically obese (ob/ob) mice. Appetite, 1987. 8(2): p. 111–23.

14. Lin, L., et al., Ghrelin receptor regulates appetite and satiety during aging in mice by regulating meal frequency and portion size but not total food intake. J Nutr, 2014. 144(9): p. 1349–55.

15. Richard, C.D., V. Tolle, and M.J. Low, Meal pattern analysis in neural-specific proopiomelanocortin-deficient mice. Eur J Pharmacol, 2011. 660(1): p. 131–8.

16. Ladenheim, E.E., et al., Disruptions in feeding and body weight control in gastrin-releasing peptide receptor deficient mice. J Endocrinol, 2002. 174(2): p. 273–81.

17. Cummings, B.P., et al., Chronic administration of the glucagon-like peptide-1 analog, liraglutide, delays the onset of diabetes and lowers triglycerides in UCD-T2DM rats. Diabetes, 2010. 59(10): p. 2653–61.

18. Horowitz, M., et al., Effect of the once-daily human GLP-1 analogue liraglutide on appetite, energy intake, energy expenditure and gastric emptying in type 2 diabetes. Diabetes Res Clin Pract, 2012. 97(2): p. 258–66.

19. Beiroa, D., et al., GLP-1 agonism stimulates brown adipose tissue thermogenesis and browning through hypothalamic AMPK. Diabetes, 2014. 63(10): p. 3346–58.

20. Larsen, P.J., et al., Systemic administration of the long-acting GLP-1 derivative NN2211 induces lasting and reversible weight loss in both normal and obese rats. Diabetes, 2001. 50(11): p. 2530–9.

21. van Can, J., et al., Effects of the once-daily GLP-1 analog liraglutide on gastric emptying, glycemic parameters, appetite and energy metabolism in obese, non-diabetic adults. Int J Obes (Lond), 2014. 38(6): p. 784–93.

22. Harder, H., et al., The effect of liraglutide, a long-acting glucagon-like peptide 1 derivative, on glycemic control, body composition, and 24-h energy expenditure in patients with type 2 diabetes. Diabetes Care, 2004. 27(8): p. 1915–21.

23. Raun, K., et al., Liraglutide, a long-acting glucagon-like peptide-1 analog, reduces body weight and food intake in obese candy-fed rats, whereas a dipeptidyl peptidase-IV inhibitor, vildagliptin, does not. Diabetes, 2007. 56(1): p. 8–15.

24. Porter, W.D., et al., Liraglutide improves hippocampal synaptic plasticity associated with increased expression of Mash1 in ob/ob mice. Int J Obes (Lond), 2013. 37(5): p. 678–84.

25. Quarenghi, M., et al., Weight Regain After Liraglutide, Semaglutide or Tirzepatide Interruption: A Narrative Review of Randomized Studies. J Clin Med, 2025. 14(11).

26. Berg, S., et al., Discontinuing glucagon-like peptide-1 receptor agonists and body habitus: A systematic review and meta-analysis. Obes Rev, 2025. 26(8): p. e13929.

27. Wilding, J.P.H., et al., Weight regain and cardiometabolic effects after withdrawal of semaglutide: The STEP 1 trial extension. Diabetes Obes Metab, 2022. 24(8): p. 1553–1564.

28. Frayn, K.N., Calculation of substrate oxidation rates in vivo from gaseous exchange. J Appl Physiol Respir Environ Exerc Physiol, 1983. 55(2): p. 628–34.

29. Rubino, D.M., et al., Effect of Weekly Subcutaneous Semaglutide vs Daily Liraglutide on Body Weight in Adults With Overweight or Obesity Without Diabetes: The STEP 8 Randomized Clinical Trial. Jama, 2022. 327(2): p. 138–150.

30. Weghuber, D., et al., Once-Weekly Semaglutide in Adolescents with Obesity. N Engl J Med, 2022. 387(24): p. 2245–2257.

31. Halawi, H., et al., Effects of liraglutide on weight, satiation, and gastric functions in obesity: a randomised, placebo-controlled pilot trial. Lancet Gastroenterol Hepatol, 2017. 2(12): p. 890–899.

32. Gabe, M.B.N., et al., Effect of oral semaglutide on energy intake, appetite, control of eating and gastric emptying in adults living with obesity: A randomized controlled trial. Diabetes Obes Metab, 2024. 26(10): p. 4480–4489.

33. Ten Kulve, J.S., et al., Liraglutide Reduces CNS Activation in Response to Visual Food Cues Only After Short-term Treatment in Patients With Type 2 Diabetes. Diabetes Care, 2016. 39(2): p. 214–21.

34. Baggio, L.L., et al., A recombinant human glucagon-like peptide (GLP)-1-albumin protein (albugon) mimics peptidergic activation of GLP-1 receptor-dependent pathways coupled with satiety, gastrointestinal motility, and glucose homeostasis. Diabetes, 2004. 53(9): p. 2492–500.

35. Flint, A., et al., The effect of glucagon-like peptide-1 on energy expenditure and substrate metabolism in humans. Int J Obes Relat Metab Disord, 2000. 24(3): p. 288–98.

36. Flint, A., et al., The effect of physiological levels of glucagon-like peptide-1 on appetite, gastric emptying, energy and substrate metabolism in obesity. Int J Obes Relat Metab Disord, 2001. 25(6): p. 781–92.

37. Näslund, E., et al., Glucagon-like peptide 1 increases the period of postprandial satiety and slows gastric emptying in obese men. Am J Clin Nutr, 1998. 68(3): p. 525–30.

38. Gibbons, C., et al., Effects of oral semaglutide on energy intake, food preference, appetite, control of eating and body weight in subjects with type 2 diabetes. Diabetes Obes Metab, 2021. 23(2): p. 581–588.

39. Friedrichsen, M., et al., The effect of semaglutide 2.4 mg once weekly on energy intake, appetite, control of eating, and gastric emptying in adults with obesity. Diabetes Obes Metab, 2021. 23(3): p. 754–762.

40. Acosta, A., et al., Quantitative gastrointestinal and psychological traits associated with obesity and response to weight-loss therapy. Gastroenterology, 2015. 148(3): p. 537–546.e4.

41. Cifuentes, L., et al., Genetic and physiological insights into satiation variability predict responses to obesity treatment. Cell Metab, 2025.

42. Ravussin, E., et al., Tirzepatide did not impact metabolic adaptation in people with obesity, but increased fat oxidation. Cell Metab, 2025. 37(5): p. 1060–1074.e4.

43. Fathi, P.A., M.B. Bales, and J.E. Ayala, Time-dependent changes in feeding behavior and energy balance associated with weight gain in mice fed obesogenic diets. Obesity (Silver Spring), 2024. 32(7): p. 1373–1388.

